# Evidence for Miocene overwater colonization in Caribbean *Cyrtognatha* spiders

**DOI:** 10.1101/372979

**Authors:** Klemen Čandek, Ingi Agnarsson, Greta Binford, Matjaž Kuntner

## Abstract

Island systems provide excellent arenas to test evolutionary hypotheses pertaining to gene flow and diversification of dispersal-limited organisms. Here we focus on an orbweaver spider genus *Cyrtognatha* (Tetragnathidae) from the Caribbean, with the aims to reconstruct its evolutionary history, describe its biogeographic history in the archipelago, and to estimate the timing and route of Caribbean colonization. Specifically, we test if *Cyrtognatha* biogeographic history is consistent with an ancient vicariant scenario (the GAARlandia landbridge hypothesis) or overwater dispersal. We reconstructed a species level phylogeny based on one mitochondrial (CO1) and one nuclear (28S) marker. We then used this topology to constrain a time-calibrated mtDNA phylogeny, for subsequent biogeographical analyses of over 100 originally sampled *Cyrtognatha* individuals. Our results suggest a monophyletic radiation of Caribbean *Cyrtognatha*, containing 11 to 14 species that are exclusively single island endemics. Our analyses refute vicariance and instead support an overwater colonization to the Caribbean in mid-Miocene. Having colonized Hispaniola first, *Cyrtognatha* subsequently dispersed to, and diversified on, the other islands of the Greater, and Lesser Antilles.

## Introduction

Island biogeography is concerned with colonization and diversification of organisms on islands, including empirical tests of evolutionary hypotheses pertaining to gene flow in dispersal-limited organisms^1,2^. Islands are geographically widespread and diverse, and vary in shapes and sizes, age and geologic origins, and show different degrees of isolation^3^. Darwin already recognized that this combination of attributes makes islands appealing objects of scientific study^4^. Modern biogeography recognizes the interplay among island histories, the specifics of their geography, and various attributes of organisms that inhabit them^5^.

Amongst island systems some of the best studied in terms of biogeographic research are Hawaii^6–8^, Galapagos^9–11^, Azores^12–14^, Canary^15–17^ and Solomon^18–20^ islands, as well as large continental fragments such as Madagascar^21–23^ and New Zealand^24–26^. However, the Caribbean island system^27–30^ is the single most ‘published’ island system in biogeography literature (Google Scholar title hits 237 compared with 195 for the second, New Zealand). The Caribbean Basin, also known as West Indies, lies in the tropical zone between South and North American continents, and to the east of the Gulf of Mexico. Combining over 700 islands, the Caribbean is considered among the world’s biodiversity hotspots^31–33^. In its most broad categorization, the Caribbean comprises three regions: (1) Greater Antilles with the largest islands of Cuba, Hispaniola (Dominican Republic and Haiti), Puerto Rico and Jamaica representing 90% of all land in Caribbean Sea; (2) Lesser Antilles with numerous smaller, mostly volcanic, islands and (3) the Lucayan platform archipelago (the Bahamas).

The Greater Antillean islands of Cuba, Hispaniola, and Puerto Rico, but not Jamaica, are parts of the old proto-Antillean arc that began its formation in the distant past, over 130 million years ago (MYA). Through Caribbean plate tectonics, the proto-Antillean arc drifted eastward until settling at its current location around 58 MYA^34,35^. Researchers disagree on the timing of the proto-Antillean arc connection with South or North America in the Cretaceous or even on the existence of such a connection. However, that distant past may have had little biological relevance for current biotas due to a catastrophic effect of the bolide that crashed into Yucatan around 65 MYA which arthropods would likely not have survived^36–38^. The emergence of the Greater Antilles as relevant biogeographic units is therefore more recent. Various studies estimate that earliest contiguous permanent dry land on the Greater Antilles has existed since the middle Eocene, approximately 40 MYA^38–42^.

Although it may be possible that the Greater Antilles have remained isolated from continental landmasses since the early Cenozoic, a hypothesized land bridge potentially existed around 35-33 MYA^38,43^. This land bridge, known as GAARlandia (Greater Antilles – Aves Ridge), is hypothesized to have connected the Greater Antilles with the South American continent for about 2 million years, due to a sea level drop and subsequent exposure of land at Aves Ridge. As a means of biotic evolution on the Greater Antilles, the GAARlandia hypothesis allows for a combination of overland dispersal and subsequent vicariance and can be tested with the help of time calibrated phylogenies and fossils. While patterns of relationships that are consistent with predictions based on GAARlandia have been found in some lineages^44–48^ it is not a good model for explaining the biogeographical history of others^27,29,49,50^.

Among the islands forming the Greater Antilles, Jamaica is a geological special case since it was originally a part of the Central American tectonic plate. Jamaica emerged as an island around 40 MYA but remained partially or fully submerged until its reemergence in mid Miocene around 15 MYA^30,51–53^, and was never part of the hypothetical GAARlandia landbridge. Consequently, Jamaica’s biota is distinct from other regions of Greater Antilles^54^.

The Lesser Antilles formed more recently. Northward of Guadalupe they split into two arches of distinct origins. The older, outer arc formed volcanically in Eocene-Oligocene, but its islands are largely composed of limestone and have undergone orogenic uplift since the Miocene. The Lesser Antilles’ inner arc is of more recent volcanic origin (<10 MYA) and its islands continue to be formed^55–59^. With no history of continental connection, most of the Lesser Antilles have been completely isolated for at least a few million years, and thus their biotas must have originated via overwater dispersal^30,60,61^.

Spiders and other arachnids are emerging as model organisms for researching biogeography of the Caribbean^27,44,48,62–64^. Spiders are globally distributed and hyperdiverse (~47,000 described of roughly 100,000 estimated species^65,66^) organisms that vary greatly in size, morphology and behavior, habitat specificity, and importantly, in their dispersal biology^67–69^. While some spiders show good active dispersal^70^, others are limited in their cursorial activities but exhibit varying passive dispersal potential. Many species are able to passively drift on air currents with behavior called ballooning^67,71^ to colonize new areas. Some genera of spiders, like *Tetragnatha* or *Nephila* are known to easily cross geographic barriers, disperse large distances, and are one of the first colonizers of newly formed islands^72–74^. These are considered to be excellent aerial dispersers, while other lineages are not as successful. For example, the primitively segmented spiders, family Liphistiidae and the mygalomorph trapdoor spiders, likely do not balloon and have highly sedentary lifestyle imposing strict limits on their dispersal potential. As a consequence, bodies of sea water or even rivers represent barriers that limit their gene flow, which leads to micro-allopatric speciation^75–80^. Unlike the above clear-cut examples, the dispersal biology of most spider lineages is unknown, and their biogeographic patterns poorly understood.

This research focuses on the tetragnathid spider genus *Cyrtognatha* and its biogeography in the Caribbean. *Cyrtognatha* is distributed from Argentina to southern Mexico and the Caribbean^65,81^. A recent revision recognized 21 species of *Cyrtognatha* but cautioned that only a fraction of its diversity is known^82^. Its biology is poorly understood as these spiders are rarely collected and studied (a single Google scholar title hit vs 187 title hits for *Tetragnatha*). Considering their phylogenetic proximity to *Tetragnatha*, as well as its described web architecture, it seems likely that *Cyrtognatha* species disperse by ballooning^83–85^. Through an intensive inventory of Caribbean arachnids, we obtained a rich original sample of *Cyrtognatha* that allows for the first reconstruction of their biogeographic history in the Caribbean. We use molecular phylogenies to reconstruct *Cyrtognatha* evolutionary history with particular reference to the Caribbean, and compare estimates of clade ages with geological history of the islands. We use this combined evidence to test the vicariant versus dispersal explanations of Caribbean colonization, and to look for a broad agreement of *Cyrtognatha* biogeographic patterns with the GAARlandia landbridge hypothesis. We also greatly expand understanding of *Cyrtognatha* diversity in the Caribbean region.

## Materials and Methods

### Sampling and identification

Material for our research was collected as a part of a large-scale Caribbean Biogeography (CarBio) project. Extensive sampling was conducted across Caribbean islands, using visual aerial search (day and night), and beating ^66,86^. Collected material was fixed in 96 % ethanol and stored at −20/−80 °C until DNA extraction. Species identification was often impossible due to juvenile individuals or lack of match with the described species (Table 1).

**Table 1:**
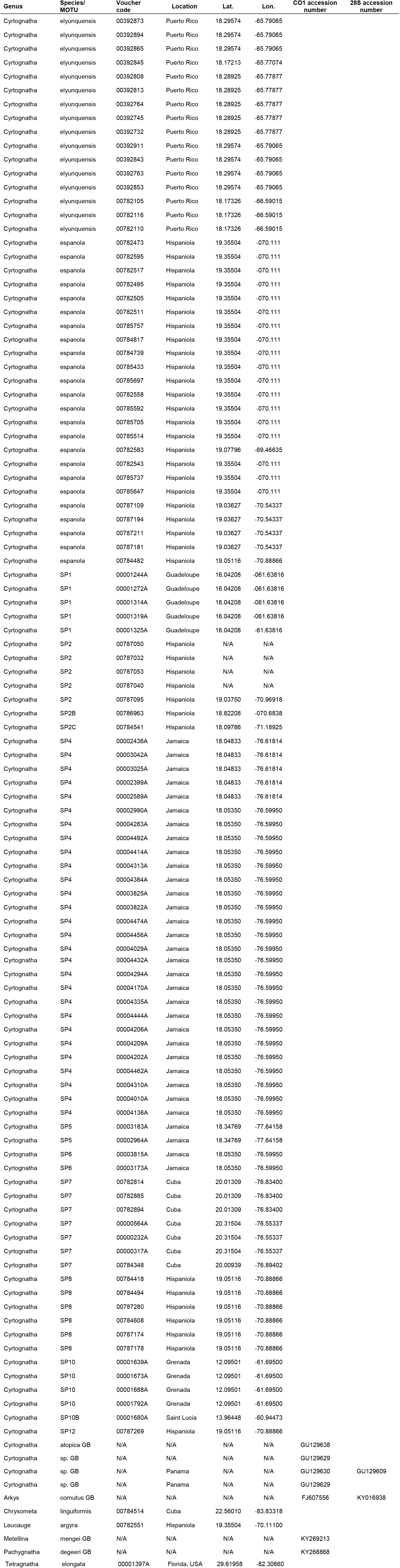
Detailed information on *Cyrtognatha* specimens and outgroups.

| Genus | Species/<br>MOTU | Voucher<br>code | Location | Lat. | Lon. | CO1 accession<br>number | 28S accession<br>number |
| --- | --- | --- | --- | --- | --- | --- | --- |
| Cyrtognatha | elyunquensis | 00392873 | Puerto Rico | 18.29574 | -65.79065 |  |  |
| Cyrtognatha | elyunquensis | 00392894 | Puerto Rico | 18.29574 | -65.79065 |  |  |
| Cyrtognatha | elyunquensis | 00392865 | Puerto Rico | 18.29574 | -65.79065 |  |  |
| Cyrtognatha | elyunquensis | 00392845 | Puerto Rico | 18.17213 | -65.77074 |  |  |
| Cyrtognatha | elyunquensis | 00392808 | Puerto Rico | 18.28925 | -65.77877 |  |  |
| Cyrtognatha | elyunquensis | 00392813 | Puerto Rico | 18.28925 | -65.77877 |  |  |
| Cyrtognatha | elyunquensis | 00392764 | Puerto Rico | 18.28925 | -65.77877 |  |  |
| Cyrtognatha | elyunquensis | 00392745 | Puerto Rico | 18.28925 | -65.77877 |  |  |
| Cyrtognatha | elyunquensis | 00392732 | Puerto Rico | 18.28925 | -65.77877 |  |  |
| Cyrtognatha | elyunquensis | 00392911 | Puerto Rico | 18.29574 | -65.79065 |  |  |
| Cyrtognatha | elyunquensis | 00392843 | Puerto Rico | 18.29574 | -65.79065 |  |  |
| Cyrtognatha | elyunquensis | 00392763 | Puerto Rico | 18.29574 | -65.79065 |  |  |
| Cyrtognatha | elyunquensis | 00392853 | Puerto Rico | 18.29574 | -65.79065 |  |  |
| Cyrtognatha | elyunquensis | 00782105 | Puerto Rico | 18.17326 | -66.59015 |  |  |
| Cyrtognatha | elyunquensis | 00782116 | Puerto Rico | 18.17326 | -66.59015 |  |  |
| Cyrtognatha | elyunquensis | 00782110 | Puerto Rico | 18.17326 | -66.59015 |  |  |
| Cyrtognatha | espanola | 00782473 | Hispaniola | 19.35504 | -070.111 |  |  |
| Cyrtognatha | espanola | 00782595 | Hispaniola | 19.35504 | -070.111 |  |  |
| Cyrtognatha | espanola | 00782517 | Hispaniola | 19.35504 | -070.111 |  |  |
| Cyrtognatha | espanola | 00782495 | Hispaniola | 19.35504 | -070.111 |  |  |
| Cyrtognatha | espanola | 00782505 | Hispaniola | 19.35504 | -070.111 |  |  |
| Cyrtognatha | espanola | 00782511 | Hispaniola | 19.35504 | -070.111 |  |  |
| Cyrtognatha | espanola | 00785757 | Hispaniola | 19.35504 | -070.111 |  |  |
| Cyrtognatha | espanola | 00784817 | Hispaniola | 19.35504 | -070.111 |  |  |

|  |  |  |  |  |  |
| --- | --- | --- | --- | --- | --- |
| Cyrtognatha | espanola | 00784739 | Hispaniola | 19.35504 | -070.111 |
| Cyrtognatha | espanola | 00785433 | Hispaniola | 19.35504 | -070.111 |
| Cyrtognatha | espanola | 00785697 | Hispaniola | 19.35504 | -070.111 |
| Cyrtognatha | espanola | 00782558 | Hispaniola | 19.35504 | -070.111 |
| Cyrtognatha | espanola | 00785592 | Hispaniola | 19.35504 | -070.111 |
| Cyrtognatha | espanola | 00785705 | Hispaniola | 19.35504 | -070.111 |
| Cyrtognatha | espanola | 00785514 | Hispaniola | 19.35504 | -070.111 |
| Cyrtognatha | espanola | 00782583 | Hispaniola | 19.07796 | -69.46635 |
| Cyrtognatha | espanola | 00782543 | Hispaniola | 19.35504 | -070.111 |
| Cyrtognatha | espanola | 00785737 | Hispaniola | 19.35504 | -070.111 |
| Cyrtognatha | espanola | 00785647 | Hispaniola | 19.35504 | -070.111 |
| Cyrtognatha | espanola | 00787109 | Hispaniola | 19.03627 | -70.54337 |
| Cyrtognatha | espanola | 00787194 | Hispaniola | 19.03627 | -70.54337 |
| Cyrtognatha | espanola | 00787211 | Hispaniola | 19.03627 | -70.54337 |
| Cyrtognatha | espanola | 00787181 | Hispaniola | 19.03627 | -70.54337 |
| Cyrtognatha | espanola | 00784482 | Hispaniola | 19.05116 | -70.88866 |
| Cyrtognatha | SP1 | 00001244A | Guadeloupe | 16.04208 | -061.63816 |
| Cyrtognatha | SP1 | 00001272A | Guadeloupe | 16.04208 | -061.63816 |
| Cyrtognatha | SP1 | 00001314A | Guadeloupe | 16.04208 | -061.63816 |
| Cyrtognatha | SP1 | 00001319A | Guadeloupe | 16.04208 | -061.63816 |
| Cyrtognatha | SP1 | 00001325A | Guadeloupe | 16.04208 | -61.63816 |
| Cyrtognatha | SP2 | 00787050 | Hispaniola | N/A | N/A |
| Cyrtognatha | SP2 | 00787032 | Hispaniola | N/A | N/A |
| Cyrtognatha | SP2 | 00787053 | Hispaniola | N/A | N/A |
| Cyrtognatha | SP2 | 00787040 | Hispaniola | N/A | N/A |
| Cyrtognatha | SP2 | 00787095 | Hispaniola | 19.03750 | -70.96918 |
| Cyrtognatha | SP2B | 00786963 | Hispaniola | 18.82208 | -070.6838 |
| Cyrtognatha | SP2C | 00784541 | Hispaniola | 18.09786 | -71.18925 |
| Cyrtognatha | SP4 | 00002436A | Jamaica | 18.04833 | -76.61814 |
| Cyrtognatha | SP4 | 00003042A | Jamaica | 18.04833 | -76.61814 |
| Cyrtognatha | SP4 | 00003025A | Jamaica | 18.04833 | -76.61814 |
| Cyrtognatha | SP4 | 00002399A | Jamaica | 18.04833 | -76.61814 |
| Cyrtognatha | SP4 | 00002589A | Jamaica | 18.04833 | -76.61814 |
| Cyrtognatha | SP4 | 00002990A | Jamaica | 18.05350 | -76.59950 |
| Cyrtognatha | SP4 | 00004283A | Jamaica | 18.05350 | -76.59950 |
| Cyrtognatha | SP4 | 00004492A | Jamaica | 18.05350 | -76.59950 |
| Cyrtognatha | SP4 | 00004414A | Jamaica | 18.05350 | -76.59950 |
| Cyrtognatha | SP4 | 00004313A | Jamaica | 18.05350 | -76.59950 |
| Cyrtognatha | SP4 | 00004384A | Jamaica | 18.05350 | -76.59950 |
| Cyrtognatha | SP4 | 00003825A | Jamaica | 18.05350 | -76.59950 |
| Cyrtognatha | SP4 | 00003822A | Jamaica | 18.05350 | -76.59950 |
| Cyrtognatha | SP4 | 00004474A | Jamaica | 18.05350 | -76.59950 |
| Cyrtognatha | SP4 | 00004456A | Jamaica | 18.05350 | -76.59950 |
| Cyrtognatha | SP4 | 00004029A | Jamaica | 18.05350 | -76.59950 |
| Cyrtognatha | SP4 | 00004432A | Jamaica | 18.05350 | -76.59950 |  |  |
| Cyrtognatha | SP4 | 00004294A | Jamaica | 18.05350 | -76.59950 |  |  |
| Cyrtognatha | SP4 | 00004170A | Jamaica | 18.05350 | -76.59950 |  |  |
| Cyrtognatha | SP4 | 00004335A | Jamaica | 18.05350 | -76.59950 |  |  |
| Cyrtognatha | SP4 | 00004444A | Jamaica | 18.05350 | -76.59950 |  |  |
| Cyrtognatha | SP4 | 00004206A | Jamaica | 18.05350 | -76.59950 |  |  |
| Cyrtognatha | SP4 | 00004209A | Jamaica | 18.05350 | -76.59950 |  |  |
| Cyrtognatha | SP4 | 00004202A | Jamaica | 18.05350 | -76.59950 |  |  |
| Cyrtognatha | SP4 | 00004462A | Jamaica | 18.05350 | -76.59950 |  |  |
| Cyrtognatha | SP4 | 00004310A | Jamaica | 18.05350 | -76.59950 |  |  |
| Cyrtognatha | SP4 | 00004010A | Jamaica | 18.05350 | -76.59950 |  |  |
| Cyrtognatha | SP4 | 00004136A | Jamaica | 18.05350 | -76.59950 |  |  |
| Cyrtognatha | SP5 | 00003183A | Jamaica | 18.34769 | -77.64158 |  |  |
| Cyrtognatha | SP5 | 00002964A | Jamaica | 18.34769 | -77.64158 |  |  |
| Cyrtognatha | SP6 | 00003815A | Jamaica | 18.05350 | -76.59950 |  |  |
| Cyrtognatha | SP6 | 00003173A | Jamaica | 18.05350 | -76.59950 |  |  |
| Cyrtognatha | SP7 | 00782814 | Cuba | 20.01309 | -76.83400 |  |  |
| Cyrtognatha | SP7 | 00782885 | Cuba | 20.01309 | -76.83400 |  |  |
| Cyrtognatha | SP7 | 00782894 | Cuba | 20.01309 | -76.83400 |  |  |
| Cyrtognatha | SP7 | 00000564A | Cuba | 20.31504 | -76.55337 |  |  |
| Cyrtognatha | SP7 | 00000232A | Cuba | 20.31504 | -76.55337 |  |  |
| Cyrtognatha | SP7 | 00000317A | Cuba | 20.31504 | -76.55337 |  |  |
| Cyrtognatha | SP7 | 00784348 | Cuba | 20.00939 | -76.89402 |  |  |
| Cyrtognatha | SP8 | 00784418 | Hispaniola | 19.05116 | -70.88866 |  |  |
| Cyrtognatha | SP8 | 00784494 | Hispaniola | 19.05116 | -70.88866 |  |  |
| Cyrtognatha | SP8 | 00787280 | Hispaniola | 19.05116 | -70.88866 |  |  |
| Cyrtognatha | SP8 | 00784608 | Hispaniola | 19.05116 | -70.88866 |  |  |
| Cyrtognatha | SP8 | 00787174 | Hispaniola | 19.05116 | -70.88866 |  |  |
| Cyrtognatha | SP8 | 00787178 | Hispaniola | 19.05116 | -70.88866 |  |  |
| Cyrtognatha | SP10 | 00001639A | Grenada | 12.09501 | -61.69500 |  |  |
| Cyrtognatha | SP10 | 00001673A | Grenada | 12.09501 | -61.69500 |  |  |
| Cyrtognatha | SP10 | 00001688A | Grenada | 12.09501 | -61.69500 |  |  |
| Cyrtognatha | SP10 | 00001792A | Grenada | 12.09501 | -61.69500 |  |  |
| Cyrtognatha | SP10B | 00001680A | Saint Lucia | 13.96448 | -60.94473 |  |  |
| Cyrtognatha | SP12 | 00787269 | Hispaniola | 19.05116 | -70.88866 |  |  |
| Cyrtognatha | atopica GB | N/A | N/A | N/A | N/A | GU129638 |  |
| Cyrtognatha | sp. GB | N/A | N/A | N/A | N/A | GU129629 |  |
| Cyrtognatha | sp. GB | N/A | Panama | N/A | N/A | GU129630 | GU129609 |
| Cyrtognatha | sp. GB | N/A | Panama | N/A | N/A | GU129629 |  |
| Arkys | cornutus GB | N/A | N/A | N/A | N/A | FJ607556 | KY016938 |
| Chrysometa | linguiformis | 00784514 | Cuba | 22.56010 | -83.83318 |  |  |
| Leucauge | argyra | 00782551 | Hispaniola | 19.35504 | -70.11100 |  |  |
| Metellina | mengei GB | N/A | N/A | N/A | N/A | KY269213 |  |
| Pachygnatha | degeeri GB | N/A | N/A | N/A | N/A | KY268868 |  |

### Molecular procedures

DNA isolation took place at University of Vermont (Vermont, USA; UVM) using QIAGEN DNeasy Tissue Kit (Qiagen, Inc., Valencia, CA), at the Smithsonian Institute in Washington, DC using an Autogenprep965 for an automated phenol chloroform extraction, and at EZ Lab (Ljubljana, Slovenia). The latter protocol involved robotic DNA extraction using Mag MAX™ Express magnetic particle processor Type 700 with DNA Multisample kit (Applied Biosystems, Foster City, CA, USA) and following modified protocols (Vidergar, Toplak & Kuntner, 2014).

We targeted two genetic markers: (1) the standard Cytochrome C oxidase subunit 1 (CO1) barcoding region, which has repeatedly been shown to be taxonomically informative in species delimitation^87,88^; and (2) the nuclear 28S gene for a subset of terminals representing all sampled species. We used the forward LCO1490 (GGTCAACAAATCATAAAGATATTGG)^89^ and the reverse C1-N-2776 (GGATAATCAGAATATCGTCGAGG)^90^ for CO1 amplification. The standard reaction volume was 25 μL containing 5 μL of Promega’s GoTaq Flexi Buffer and 0.15 μL of GoTaq Flexi Polymerase, 0.5 μL dNTP’s (2 mM each, Biotools), 2.3 μL MgCl_2_ (25 mM, Promega), 0.5 μL of each primer (20 μM), 0.15 μL BSA (10 mg/mL; Promega), 2 μL DNA template and the rest was sterile distilled water. We used the following PCR cycling protocol: an initial denaturation step of 5 min at 94 °C followed by 20 touch-up method cycles of 60 s at 94°C, 90 s at 44 °→ 54 °C, 1 min at 72 °C, followed by 15 cycles of 90 s at 94 °C, 90 s at 53.5 °C, 60 s at 72 °C and the final extension period of 7 min at 72 °C.

The primer pair for 28S were the forward 28Sa (also known as 28S-D3A; GACCCGTCTTGAAACACGGA)^91^ and the reverse 28S-rD5b (CCACAGCGCCAGTTCTGCTTAC)^92^. The standard reaction volume was 35 μL containing 7.1 μL of Promega’s GoTaq Flexi Buffer and 0.2 μL of GoTaq Flexi Polymerase, 2.9 μL dNTP’s (2 mM each, Biotools), 3.2 μL MgCl2 (25 mM, Promega), 0.7 μL of each primer (20 μM), 0.2 μL BSA (10 mg/mL; Promega), 1 μL DNA template and the rest was sterile distilled water. We used the following PCR cycling protocol: an initial denaturation step of 7 min at 96 °C followed by 20 touch-down method cycles of 45 s at 96 °C, 45 s at 62 ° C → 52 °C, 60 s at 72 °C, followed by 15 cycles of 45 s at 96 °C, 45 s at 52 °C, 60 s at 72 °C and the final extension period of 10 min at 72 °C. The PCR products were purified and sequenced at Macrogen (Amsterdam, NL).

We used Geneious v. 5.6.7^93^ for sequence assembly, editing and proofreading. For alignment, we used the default settings and the automatic optimization option in the online version of MAFFT^94^. We concatenated the CO1 and 28S matrices in Mesquite^95^.

We obtained 103 original *Cyrtognatha* CO1 sequences and mined four additional *Cyrtognatha* CO1 sequences from GenBank (Table 1). Moreover, we added three CO1 sequences from GenBank (*Arkys cornutus, Metellina mengei, Pachygnatha degeeri*) and three original CO1 sequences (*Leucauge argyra, Chrysometa linguiformis, Tetragnatha elongata*) to be used as outgroups. We obtained 22 original sequences of 28S gene fragment representing all putative species of *Cyrtognatha* and included one from GenBank. Additionally, we incorporated three original 28S sequences (*Leucauge argyra, Chrysometa linguiformis, Tetragnatha elongata*) and a single one from GenBank (*Arkys cornutus*) to be used as outgroups (Table 1). The concatenated matrix contained 1244 nucleotides (663 for CO1 and 581 for 28S).

### Species delimitation

Because the current taxonomy of *Cyrtognatha* based on morphology is highly incomplete^82^, we undertook species delimitation using CO1 data. To determine the molecular taxonomic operational units (MOTUs), we used four different species delimitation methods, each with its online application: PTP (Poisson tree process)^96^, mPTP (multi-rate Poisson tree process)^97^, GMYC (generalized mixed yule coalescent)^98^ and ABGD (automatic barcode gap discovery)^99^. We ran these species delimitation analyses using the default settings, with the input tree for GYMC from BEAST2^100^ and the input trees for PTP and mPTP from MEGA 6.0^101^.

### Phylogenetic analyses

We used MrBayes^102^ to reconstruct an all-terminal phylogeny for a complete set of our original *Cyrtognatha* material and outgroups using CO1 (Table 1). For Bayesian analysis we used the Generalised time-reversible model with gamma distribution and invariant sites (GTR+G+I) as suggested by AIC and BIC criterion in jModelTest2^103^. We ran two independent runs, each with four MCMC chains, for 100 million generations, with a sampling frequency of 1000 and relative burn-in set to 25%. The starting tree was random.

For a species level phylogeny, we then selected two individuals per MOTU and added 28S sequence data for two partitions and analyzed this concatenated dataset under a Bayesian framework. As above, jModelTest2 suggested GTR+G+I as the appropriate model, this time for both partitions. These analyses had 28 terminals including outgroups (Table 1). The settings in MrBayes were as above, but the number of MCMC generations was set to 30 million. Due to high mutation rates in noncoding parts of nuclear genes like 28S, insertions and deletions accumulate through evolution^104,105^, resulting in numerous gaps in a sequence alignment. We treated gaps as missing data but also ran additional analyses applying simple gap coding with FastGap^106^.

### Molecular dating analyses

We used BEAST2^100^ for time calibrated phylogeny reconstruction (chronogram). We used a single CO1 sequence per MOTU and trimmed the sequences to approximately equal lengths. We then modified the xml file in BEAUti^100^ to run three different analyses. The first analysis was run using GTR+G+I as suggested by jModelTest2. The second analysis employed the package and model bModelTest^107^. The third analysis used the package and RBS model^108^. All parameters were set to be estimated by BEAST. We calibrated a relaxed log normal clock following Bidegaray-Batista and Arnedo^109^ with the ucld.mean set as normal distribution with mean value of 0.0112 and standard deviation of 0.001, and the ucld.stdev set to exponential distribution with the mean of 0.666. We ran an additional analysis using a fossil calibration point on the basal node of Caribbean *Cyrtognatha* clade. *Cyrtognatha weitschati,* known from Dominican amber of Hispaniola, is hypothesized to be 13.65 – 20.41 million years old. We used an exponential prior with 95 % confidence interval spanning from a hard lower bound at 13.65 MYA to the soft upper bound at 41 MYA. This upper bound corresponds with the time of Hispaniola appearance. We selected Yule process as a tree prior and constrained the entire topology for BEAST analyses based on the results from the above described species level phylogeny. The trees were summarized with TreeAnnotator^100^, with 20% burn-in based on a Tracer^110^ analysis, target tree set as Maximum clade credibility tree and node heights as median heights.

All metafiles from BEAST and MrBayes were evaluated in Tracer to determine burn-in, to examine ESS values and to check for chain convergence. For visualization of trees we used FigTree^111^. All MrBayes and BEAST analyses were run on CIPRES portal^112^.

### Ancestral area estimation

We used BioGeoBEARS^113^ in R version 3.5.0^114^ to estimate ancestral range of *Cyrtognatha* in the Caribbean. We used a BEAST produced ultrametric tree from the above described molecular dating analysis as an input. We removed the outgroup *Tetragnatha elongata* and conducted the analyses with the 13 *Cyrtognatha* MOTUs from six areas (Hispaniola, Jamaica, Puerto Rico, Cuba, Lesser Antilles and Panama). We estimated the ancestral range of species with all models implemented in BioGeoBEARS: DEC (+J), DIVALIKE (+J) and BAYAREALIKE (+J). We used log-likelihoods (LnL) with Akaike information criterion (AIC) and sample-size corrected AIC (AICc) scores to test each model’s suitability for our data. All of our *Cyrtognatha’s* MOTUs are single island endemics, therefore we were able to reduce the parameter “max_range_size” to two^29,115,116^.

## Results

We collected 103 *Cyrtognatha* individuals from Cuba, Jamaica, Dominican Republic/Hispaniola, Puerto Rico and Lesser Antilles (Fig. 1, Table 1). We confirmed that all individuals are morphologically *Cyrtognatha*. Most species are undescribed but we identified two known species: *C. espanola* (Bryant, 1945) and *C. elyunquensis* (Petrunkevitch, 1930) comb. nov., previously placed in *Tetragnatha*^65^.

**Figure 1:**
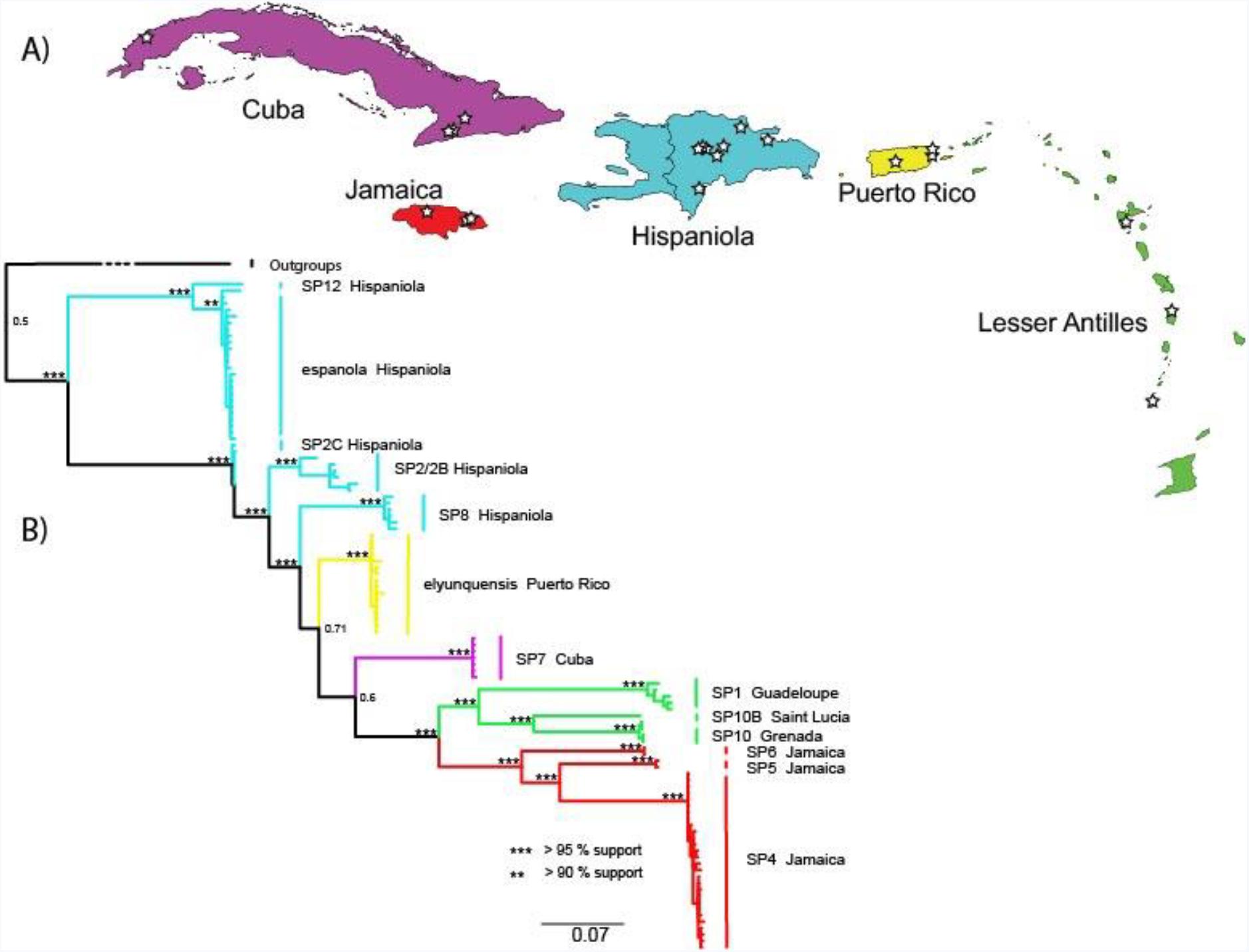
A) Map of the Caribbean with indicated sampling localities. B) The all-terminal mitochondrial Bayesian phylogeny of *Cyrtognatha*. Branch colors match those of the islands in A. Notice that all putative species form exclusively single island endemic pattern.

We obtained CO1 sequences for all *Cyrtognatha* individuals. Using computational methods for species delimitation our *Cyrtognatha* dataset is estimated to contain from 11 to 14 MOTUs (Supplementary Note S1). The results from PTP, mPTP and ABGD were mostly consistent, disagreeing only on the status of three putative species. To these species that are supported by some but not all analyses, we added the label B or C after the species name: *Cyrtognatha* SP10B, *Cyrtognatha* SP2B and *Cyrtognatha* SP2C. On the other hand, we dismiss the results from GMYC analyses using either a single versus multiple threshold option, which failed to recover reliable MOTUs. The composition of our dataset is most likely not compatible with GMYC method as it cannot detect switches between inter-and intraspecific branching patterns, offering us from 1 to 39 MOTUs.

The two gene and the CO1 phylogenies yielded nearly identical networks for the Caribbean taxa, but strikingly different phylogenies due to different root placement (Fig. 2, Supplementary Fig. S1). Therefore, we ran an additional analysis constraining the root of the mtDNA phylogeny to reflect that of the two gene phylogeny, with otherwise the same settings (Fig. 1, Supplementary Fig. S2). Our unconstrained all-terminal Bayesian phylogeny supports Caribbean *Cyrtognatha* monophyly, albeit with only three non-Caribbean samples. Most terminal clades were well supported with lower supports for some deeper nodes (Supplementary Fig. S1). This phylogeny strongly recovers all putative species groups as single island endemics. Furthermore, all geographic areas harbor monophyletic lineages, with the exception of Hispaniola that supports two independent clades. The unconstrained all-terminal CO1 phylogeny recovers the Lesser Antillean clade as sister to all other Caribbean taxa. However, this relationship is not recapitulated in the concatenated, species level, phylogeny (Fig. 2, Supplementary Fig. S3). The concatenated phylogeny also supports monophyly of the Caribbean taxa but recovers the clade of *C. espanola* and *C*. SP12 from Hispaniola as sister to all other Caribbean *Cyrtognatha.* The species level phylogeny is generally better supported, with the exception of a clade uniting species from Lesser Antilles, Cuba, Hispaniola and Puerto Rico. In both Bayesian analyses the chains successfully converged and ESS as well as PRSF values of summarized MCMC runs parameters were appropriate^102^.

**Figure 2:**
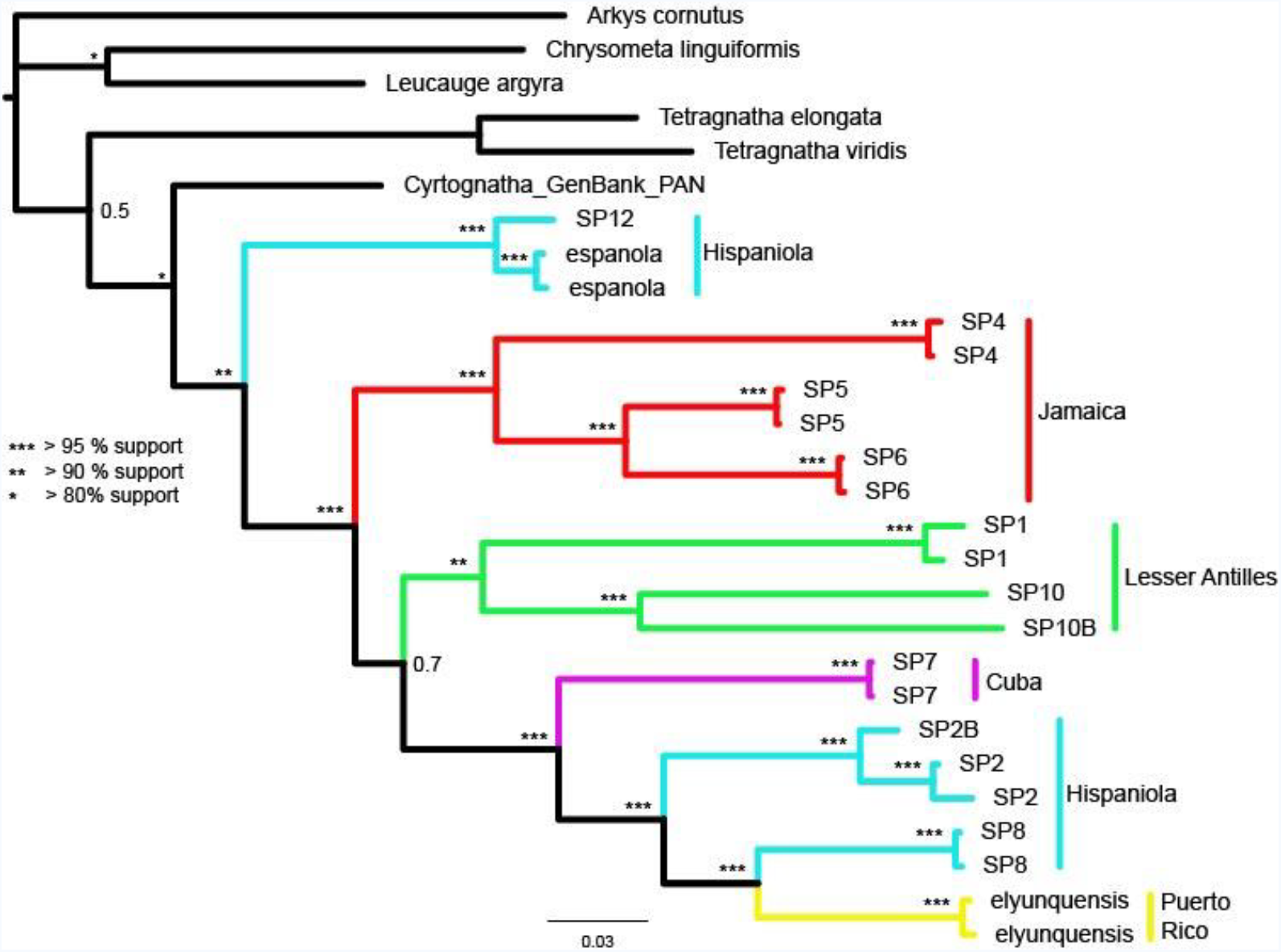
Species level Bayesian phylogeny of *Cyrtognatha* based on CO1 and 28S. Relationships agree with *Cyrtognatha* and Caribbean *Cyrtognatha* monophyly.

Chronograms produced by BEAST, using either exclusively CO1 mutation rate or incorporating the additional fossil for time calibration, exhibited very similar time estimates (Fig. 3, Supplementary Fig. S5). We decided to proceed with the mutation rate-only calibrated phylogeny for further analyses because it is less likely to contain known potential biases when calibrating with scarcely available fossils and geological information^117,118^. The molecular dating analyses based on the three different models in BEAST largely agreed on node ages with less than 1 million years variation. However, the log files from the chronogram based on GTR+G+I model consistently exhibited low ESS values (<50), even with MCMC number of generations having been increased to 200 million. The analyses using the remaining models, RBS and bModelTest, were more appropriate since MCMC chains successfully converged, and the lowest ESS values were 981 and 2214 respectively, thus far exceeding the suggested 200. Additional examination of the log files produced by bModelTest phylogeny with bModelAnalyzer from AppStore.exe^107^ revealed that MCMC chains spent most time in modified TN93 model with the code 123143 which contributed for 49.56% of posterior probability (for details on bModelTest method of model selection see ^107^). The BEAST chronogram using bModelTest (Fig. 3, Supplementary Fig. S4) yielded the best supported results, amongst the above mentioned approaches, based on ESS values, and was therefore used in subsequent biogeographical analyses. This chronogram (Fig. 3) supports a scenario in which *Cyrtognatha* diverged from its sister genus *Tetragnatha* at 18.7 MYA (95% HDP: 12.8 – 26.7 MYA). The Caribbean clade is estimated to have split from the mainland *Cyrtognatha* (represented here by a species from Panama) 15.0 MYA (95% HDP: 10.5 – 20.7 MYA). The clade with lineages represented on Lesser Antilles diverged from those on Greater Antilles at 11.5 MYA (95% HDP: 8.3 – 15.6 MYA).

**Figure 3:**
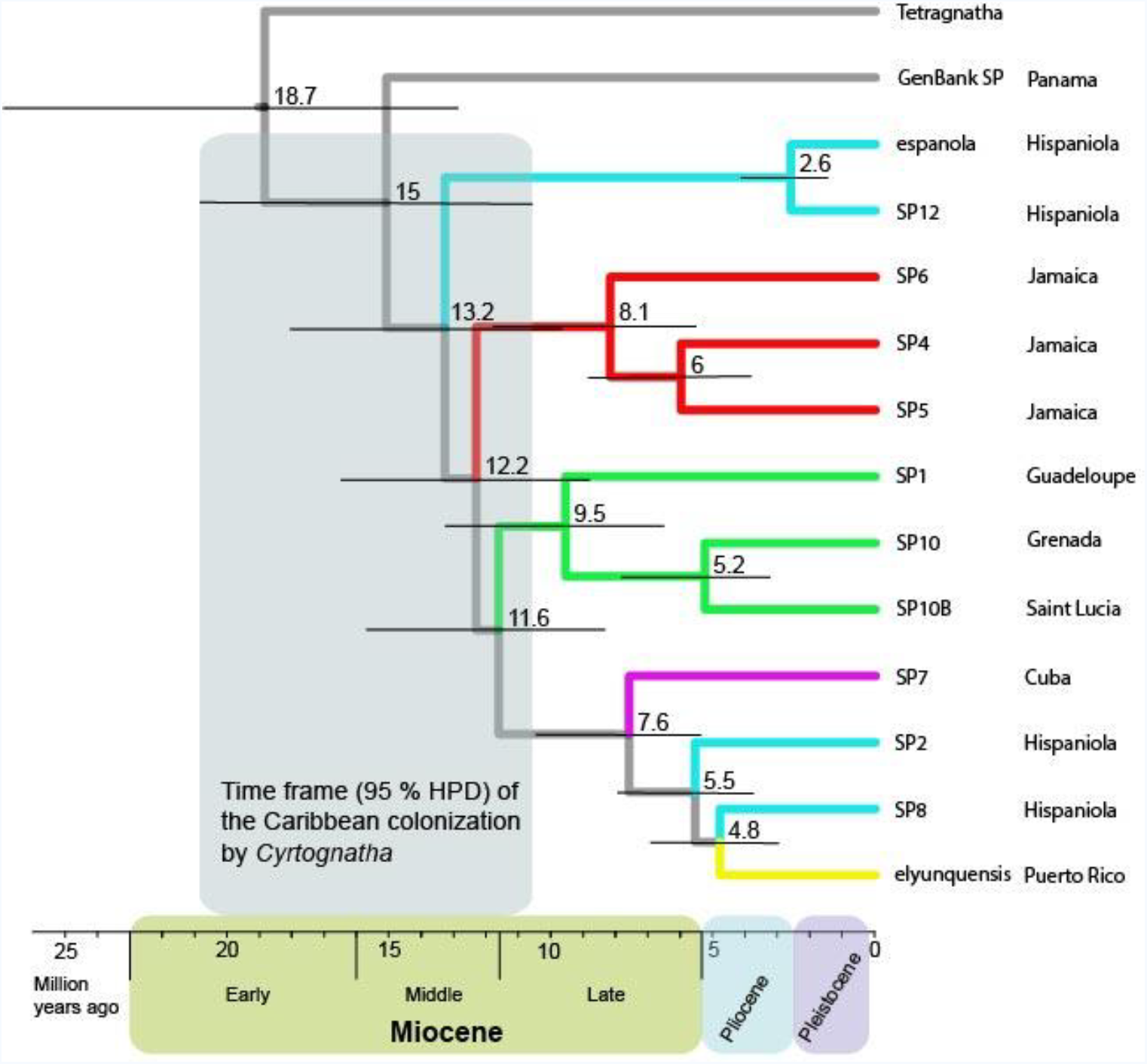
Time-calibrated BEAST phylogeny of *Cyrtognatha.* This chronogram suggests *Cyrtognatha* colonized the Caribbean in mid-Miocene and refutes ancient vicariant scenarios. The lack of any land bridge connection of the Caribbean with mainland at least since early Oligocene (cca. 33 MYA; GAARlandia) suggests that colonization happened by overwater dispersal. Confidence intervals of clade ages agree with geological history of Caribbean islands.

The comparison of all six models of ancestral area estimation with BioGeoBEARS recovered DIVALIKE+J as most suitable for our data due to highest LnL scores in both AIC and AICc tests (Supplementary Table S1). The estimation of ancestral states suggests that the most recent common ancestor of all Caribbean *Cyrtognatha* in our dataset most likely (62%) resided on Hispaniola (Fig. 4). Moreover, all the Greater Antillean island clades as well as the Lesser Antillean clade most likely originated from Hispaniola with the following probability: Jamaican clade (47%), Lesser Antillean clade (40%), Cuban clade (63%) and Puerto Rican clade (89%) (Fig. 4, Supplementary Table S1).

**Figure 4:**
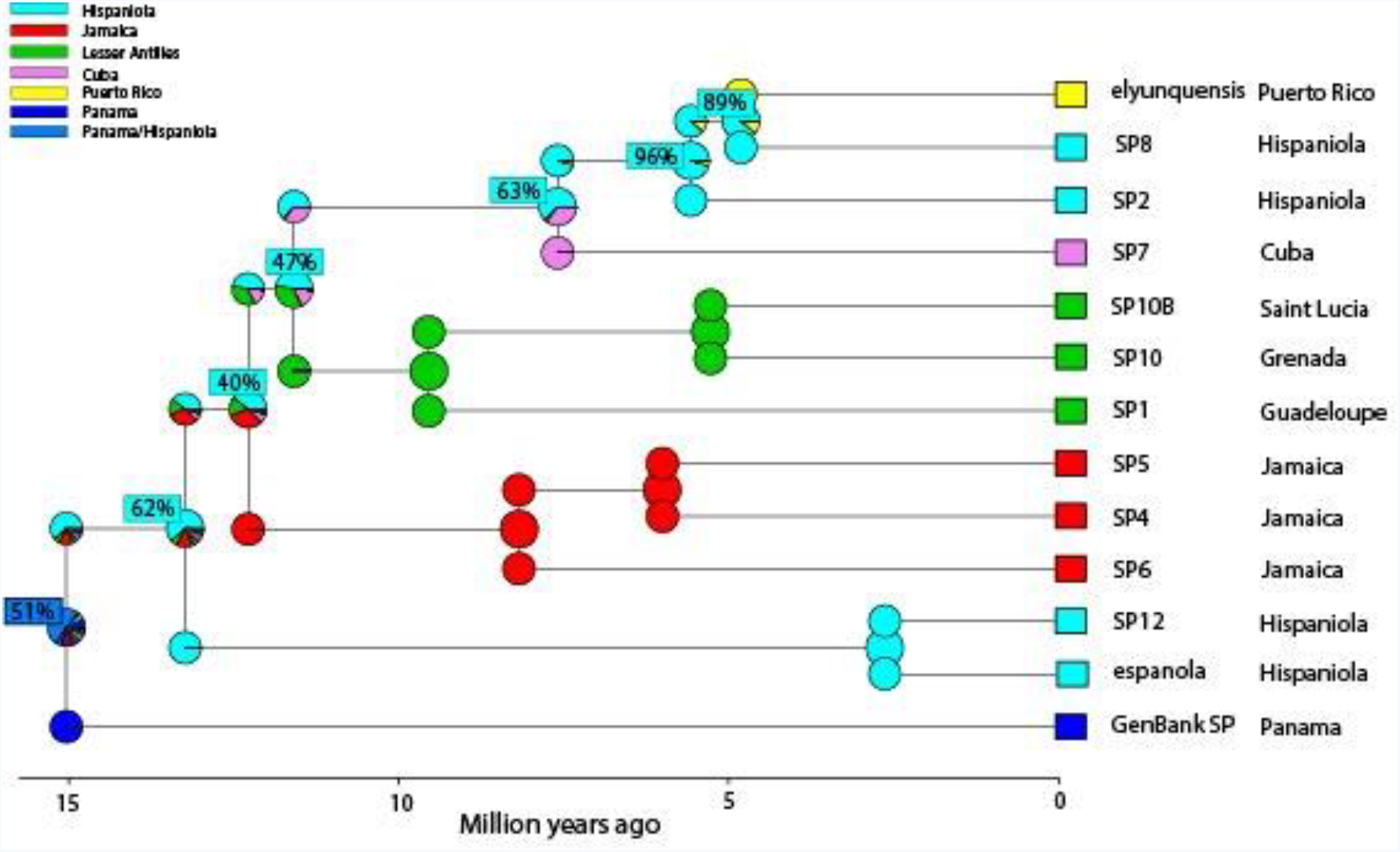
Ancestral area estimation of *Cyrtognatha* with BioGeoBEARS. The biogeographical analysis, using the most suitable model for our data (DIVALIKE + J, max_range_size = 2), revealed that Hispaniola was most likely colonized first. Following colonization, *Cyrtognatha* diversified within Hispaniola and subsequently dispersed from there to all other islands of the Caribbean.

## Discussion

We reconstruct the first *Cyrtognatha* phylogeny using molecular data from over 100 individuals of this rarely collected group. Our results support *Cyrtognatha* as a relatively young clade, having diverged from a common ancestor with its sister genus *Tetragnatha*, in early-to mid-Miocene, and colonized the Caribbean in mid-Miocene. As we discuss below, these estimated ages, combined with the phylogenetic patterns, refute vicariant explanations of their Caribbean origin, including the GAARlandia hypothesis. Instead, the patterns suggest colonization of Hispaniola, and subsequent dispersal to other islands.

The all-terminal phylogeny (Fig. 1) reveals clear patterns of exclusively single island endemic (SIE) species. This holds true even for the three MOTUs on the Lesser Antilles island group, as they appear on Guadelupe, St. Lucia and Grenada (Table 1). Even in the absence of the oceanic barriers, i.e. within the larger islands, we find evidence of short range endemism^119^. While we do not claim to have thorough regional sampling, we find patterns of local endemism in regions where our sampling is particularly dense, providing the strongest test with available data. Many Caribbean spiders such as *Spintharus*^44^, *Micrathena*^120^ *Selenops*^121^ and *Nops*^122^, as well as other arachnid lineages such as *Amblypygi*^64^ and *Pseudoscorpions*^63^, demonstrate a similar pattern. The distribution and quantity of SIEs depends on island properties such as maximum elevation, size, isolation and geological age^123–127^. While our focus was not on the effect of physical properties of islands on SIEs, the patterns seem to point towards a higher number of SIEs on the islands with a higher maximum elevation: Hispaniola (3098 m) is occupied by 4 or 6 MOTUs (depending on the delimitation method), Jamaica (2256 m) by three MOTUs and all other islands (< 2000 m) by a single MOTU. The Caribbean islands, with the exception of Hispaniola, also harbor exclusively monophyletic *Cyrtognatha* lineages. We might explain this observed pattern with a combination of the niche preemption concept and organisms’ dispersal ability^128–130^. A combination of the first colonizer’s advantageous position to occupy empty niches and rare overwater dispersal events of their closely related species leads to competitive exclusion and lower probability for newcomers to establish viable populations on already occupied islands^131–133^. While niche preemption is better studied in plants, it is also applicable to animals, including spiders^1,134^.

*Cyrtognatha* most likely colonized the Caribbean through long distance overwater dispersal. An ancient vicariant hypothesis would predict that the early proto-Antilles were connected to the continental America and were colonized in the distant past, possibly over 70 MYA^135^ and the GAARlandia landbridge putatively existed around 35-33 MYA. These hypothetical scenarios are clearly not reflected in our BEAST chronogram (Fig. 3) in which we estimate that the Caribbean *Cyrtognatha* split from its continental population as late as 13.2 MYA. This suggests that the genus *Cyrtognatha* is much younger than the most reasonable possible vicariant timeframe. Likewise, the reconstructed biogeographic patterns fail to support vicariant scenarios. First, the Jamaican lineage split from Hispaniola soon after colonization (12.2 MYA) even though Jamaica was never a part of the proto-Antilles. Secondly, Puerto Rico was a part of the proto-Antilles but was colonized only recently (4.8 MYA). The results of Jamaican and Puerto Rican colonization from Hispaniola thus are most consistent with a scenario of colonization by overwater dispersal.

The mid-Miocene (ca. 15 MYA) is considered as the start of the modern Earth^136^ in that the climate began to stabilize and the ocean currents started to take their current form. This combination of events enabled the colonization of the Caribbean islands from eastern-northern parts of South America for example via vegetation rafts passively drifting with water currents^137^. That also meant that the wind directions and the hurricane paths most likely resembled those of today^138^, from East to West direction^139^. In fact, hurricanes may create numerous dispersal/colonization opportunities, especially for the organisms with poor active dispersal abilities^135,140^. Wind directions and tropical storms are relevant for tetragnathid spiders like *Cyrtognatha* that disperse by ballooning and could facilitate their colonization of the Caribbean islands in a stepping stone^141^ or leap-frog^142^ manner.

With the examination of the relationships in the time calibrated phylogeny (Fig. 3), we predict that a single colonization of the Caribbean from the continental America occurred sometime between 10.5 and 20.7 MYA. The most likely scenario indicates the original colonization of the Greater Antilles (Hispaniola; Fig. 4). Such patterns of colonization of Greater Antilles in Miocene are also evident in many other lineages including vertebrates, invertebrates and plants ^29,41,143–150^. A more rigorous test of number and directionality of colonization pathways would require more thorough sampling across potential source populations on the mainland.

Our inference of ancestral ranges proposes an early within island diversification of *Cyrtognatha* ancestors occupying Hispaniola and predict that Hispaniola is the ancestral area for all Caribbean clades (Fig. 4, Supplementary Fig. S5). The path of colonization does not resemble a straightforward pattern such as the stepping-stone pattern. The colonization sequence seems more random or resembles a “leap-frog” pattern. In our case the clear example of island being “leap frogged” is Puerto Rico. A leap frog pattern could indicate a role of hurricanes in movement among Caribbean islands^135,151^.

## Conclusions

Our phylogenetic analysis of the tetragnathid spider genus *Cyrtognatha* facilitates reconstruction of its biogeographic history in the Caribbean. The ancestor of this relatively young lineage appears to have colonized the Caribbean overwater in the Miocene and further diversified into an exclusively single island endemic biogeographic pattern seen today. Further sampling of these rarely collected spiders in continental America is needed to confirm the timing, number and source of colonization of the Caribbean and to contrast those from other Caribbean spider clades. For example, *Spintharus*^44,48^ and *Deinopis* (Chamberland et al. ‘in review’) patterns clearly support ancient vicariance, but *Argiope*^27^ readily disperses among the islands. *Tetragnatha*, the sister lineage of *Cyrtognatha*, may prove to be of particular interest in comparison, because its biogeographic history on the islands may mirror their global tendency towards repeated colonization of even most remote islands.

## Contributions

Research design: K.Č., I.A., G.B., M.K. Material acquisition and molecular procedures: I.A., G.B., K.Č., M.K. Data analyses: K.Č. K.Č. wrote the first draft of the paper. All authors contributed to writing and revising the paper.

## Data availability

All data generated in this study and protocols needed to replicate it are included in this published article and its Supplementary material files.

## Acknowledgements

We thank the entire CarBio team (http://www.islandbiogeography.org/participants.html) for collecting the material across the Caribbean. Moreover, we thank Lisa Chamberland and other members of the Agnarsson lab (http://www.theridiidae.com/lab-members.html) for the help with material sorting and molecular procedures. This work was supported by grants from the National Science Foundation (DEB-1314749, DEB-1050253), and the Slovenian Research Agency (J1-6729, P1-0236, BI-US/17-18-011).

